# Cylindrical, pylon-like structures with helix recesses enhance coral larval recruitment

**DOI:** 10.1101/2025.10.06.680805

**Authors:** Jessica Reichert, Hendrikje Jorissen, Marina E. Rottmueller, Allison D. Nims, Lomani H. Rova, Crawford Drury, the R3D Consortium, Joshua S. Madin

## Abstract

The decline of coral reefs requires scalable restoration strategies to enhance natural recovery processes such as coral larval recruitment. Previous research has shown that conical structures with helix recesses substantially increase settlement and early survival. However, the applicability of this microhabitat design with helix recesses in broader engineering contexts has yet to be assessed. Hereby, we tested (1) whether helix recesses can be transferred from conical dome geometries to space-efficient cylindrical pylon-like geometries, and (2) whether they can be implemented with different materials. In a field experiment in Kāneʻohe Bay, Hawaiʻi, coral recruitment was monitored over six months on cylindrical and conical structures incorporating an optimized helix profile. Cylindrical modules supported recruitment densities similar to those on conical designs, demonstrating a successful transfer of the microhabitat design to compact geometries. Planar recruit densities were ∼300 times higher across all structures, compared to those observed on nearby natural reefs. These results show that the helix recess design is functionally robust across both module shapes and materials. Cylindrical structures with integrated helix recesses, therefore, represent a practical, low-cost design element that can be incorporated in coastal engineering and restoration projects to enhance coral settlement.

**Graphical abstract:** 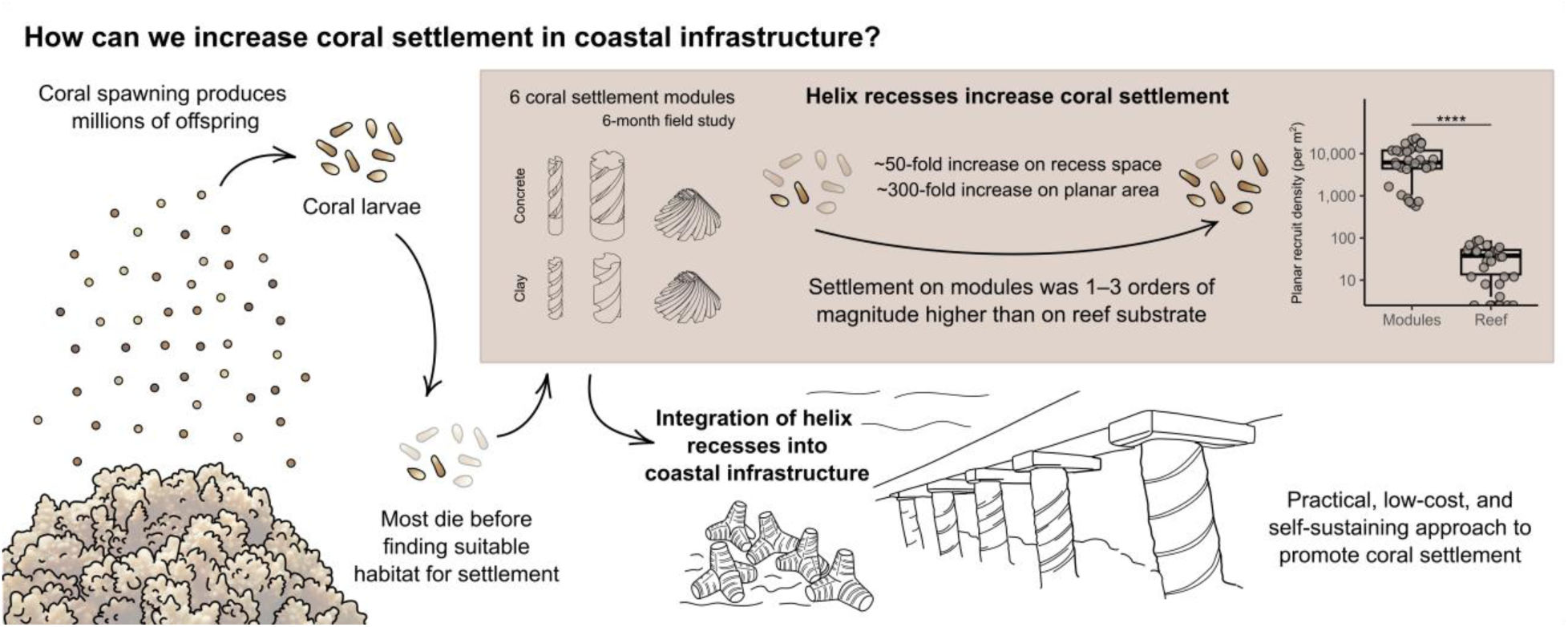

## 1. Introduction

Coral reefs are degrading worldwide due to the impacts of climate change and local stressors (Hughes et al., 2017). Lowering global carbon emissions is our best way to combat this trend globally, yet more effective and scalable restoration strategies are needed to address this trend locally (Edwards et al., 2024; Hughes et al., 2023). Scaling up restoration by harnessing the natural capacity of corals to produce millions of sexually derived larvae presents a promising approach (Boström-Einarsson et al., 2020; Madin et al., 2025). The transition from planktonic larvae to settled polyps is a crucial bottleneck in the coral life cycle, characterized by high mortality and a strong dependence on suitable settlement habitats (Martinez and Abelson, 2013; Nozawa and Harrison, 2008). Designing artificial structures that provide optimal settlement microhabitats can therefore substantially enhance recruitment and restoration success.

Coral larval settlement and survival depend on interacting physical, chemical, and biological factors, including light, substrate properties, hydrodynamics, settlement cues, predation, and competition. Light intensity and turbidity influence recruit photosynthesis, algal fouling, and the composition of CCA, green, and brown algae, which in turn affects recruit survival (Ramsby et al., 2024). Coral larvae typically settle in cryptic, low-light habitats (<500 µmol photons m⁻² d⁻¹), yet survival is reduced under very low light conditions (<100 µmol photons m⁻² d⁻¹), (Brunner et al., 2022). Water flow and hydrodynamics shape larval dispersal, transport, and settlement. Coral larvae, which are poor swimmers, often settle in millimeter-scale ridges, where microtopography creates low-flow microenvironments that support settlement (Hata et al., 2017; Levenstein et al., 2022a). Therefore, habitat shape and surface topography strongly influence larval settlement and survival. Coral larvae commonly settle in cryptic microhabitats such as crevices (Brunner et al., 2022; Mundy and Babcock, 1998). Crevice size and geometry (aperture and depth) are key factors influencing coral recruitment (Doropoulos et al., 2016; Nozawa, 2008). Different coral taxa exhibit distinct settlement preferences and survival rates across depths and microhabitats, including exposed, vertical, and cryptic surfaces (Edmunds et al., 2004). In addition, surface microtopography strongly influences larval settlement: coral larvae often prefer recess-shaped microtopographies, but preferences differ among species (Nozawa, 2008; Petersen et al., 2005). Further, porous surfaces with specific pore sizes and morphologies provide additional settlement microhabitats for coral larvae (Hoog Antink et al., 2018). Specific engineered surface microtopographies, such as micro-sized sine wave recesses or the Sharklet AF™ pattern, can also reduce biofouling (Erramilli and Genzer, 2019; Myan et al., 2013). Biological cues also play a crucial role in coral larval settlement. For example, the presence of a biofilm is essential for coral larval settlement (Petersen et al., 2005). Also, CCA are well-known inducers of coral larval settlement (Harrington et al., 2004). Their chemical compounds, such as glycoglycerolipids and betaine lipids, and associated bacterial communities, act as settlement cues for coral larvae (Gómez-Lemos et al., 2018; Jorissen et al., 2021; Tebben et al., 2011). The material of the settlement substrate can influence both coral recruitment and the development of early benthic communities (Levenstein et al., 2022b). However, site-specific factors often outweigh substrate material in shaping benthic communities and coral recruitment (Burt et al., 2009). The presence of other early benthic communities, which involve competition, such as bryozoans, can negatively impact post-settlement survival of corals (Doropoulos et al., 2016). Predation, although difficult to quantify directly in the field, is considered the primary reason for post-settlement mortality (Cooper et al., 2014). Grazers may feed indiscriminately on recruits, especially on exposed surfaces, while corallivores and cryptic predators also target recruits in sheltered areas (Brandl et al., 2014, Doropoulos et al., 2016). Protective microhabitats are therefore crucial for recruit survival.

Given the importance of reef structural complexity for ecosystem function, current restoration approaches increasingly use 3D printing to create artificial reefs from materials such as ceramic, cement, and geopolymer mortars (Hoog Antink et al., 2018; Yoris-Nobile et al., 2023). 3D printing enables the fabrication of structures that are tailored to restoration goals and mimic the complex architecture of natural reefs (Berman et al., 2023; Chamberland et al., 2017; Levy et al., 2022; Perricone et al., 2023). Large-scale reef-mimicking modules include MARS units by Reef Design Lab, Ørsted, and WWF’s limestone reef blocks in the Kattegat Sea, 3D-printed concrete structures in Florida and Louisiana, and Reef Balls. At smaller scales, 3D-printed substrates for coral settlement include SECORE’s ceramic seeding units and Archireef’s terracotta tiles.

In line with these developments, our previous work identified optimal settlement microhabitats using 3D-printed clay modules deployed in Kāneʻohe Bay, Hawaiʻi (Reichert et al., 2024). We tested seven conical designs incorporating different surface features (holes, pockets, and recesses) across a range of sizes. We found that specific helix recesses dramatically enhanced settlement (by ∼80-fold) and post-settlement survival (by 20-50-fold over one year) of the dominant spawning coral *Montipora capitata*, compared to control domes lacking these features. The most successful ‘superdome’ design incorporated 12 narrow recesses that created microhabitats with low light levels (∼50-60 µmol m⁻² s⁻¹) where recruits preferentially settled along cryptic inner edges. While conical domes provided an effective experimental platform, their shape limits direct application in coastal engineering, where structures are often cylindrical, modular, or pylon-like. A central, unresolved question is whether helix recesses represent a generalizable microhabitat principle rather than a geometry-specific solution. Because cylindrical and pylon-like geometries underpin many coastal protection systems, such as pilings, seawalls, and breakwaters, establishing whether helix recesses function on these infrastructure-relevant forms is critical for their integration into real-world coastal engineering and hybrid reef solutions (Higgins et al., 2022). Transferring recess designs from cones to cylinders is challenging, because macro-geometry affects flow separation, shear, and light regimes, all of which influence larval contact and settlement (Koehl, 2007; Levenstein et al., 2022a). Additionally, previous work focused exclusively on fired clay, leaving open whether the same recess design functions when implemented in concrete, the dominant and most scalable material for coastal infrastructure.

To address these gaps, this study evaluates whether helix recesses function as a generalizable microhabitat design across geometries, materials, and recess dimensions. First, we test whether the optimized helix microhabitat can be successfully transferred across macro-geometries by applying it to space-efficient, pylon-like cylindrical rods and comparing their performance with that of conical superdomes deployed simultaneously. Second, we assess whether helix recesses are material-robust by evaluating coral recruitment on modules constructed from cast concrete versus 3D-printed clay. Third, we leveraged variation in recess dimensions arising from 3D-printing constraints to examine whether recess dimensions generate functional trade-offs between early settlement and longer-term recruit persistence. Together, these analyses will elucidate whether helix recesses constitute a scalable, engineering-ready design principle suitable for integration into hybrid coastal infrastructure.

## 2. Materials and Methods

### 2.1 Experimental design

We conducted a field experiment to evaluate the suitability of helix recess designs on rod-shaped structures. The experiment compared coral recruitment and survival of six module geometries of settlement modules: four cylindrical rod designs (‘small clay rod’, ‘large clay rod’, ‘small concrete rod’, and ‘large concrete rod’) and two conical superdome designs (‘superdome clay’ and ‘superdome concrete’), (Figure 1). Settlement modules (n = 6 per cylindrical rod design, n = 4 per superdome clay, and n = 3 per superdome concrete) were deployed at Reef 13 in Kāneʻohe Bay, Oʻahu, Hawaiʻi, one week prior to the predicted *Montipora capitata* mass spawning in June 2024. Recruitment was monitored at 1 week, 2 months, and 6 months post-spawning.

**Figure 1:**
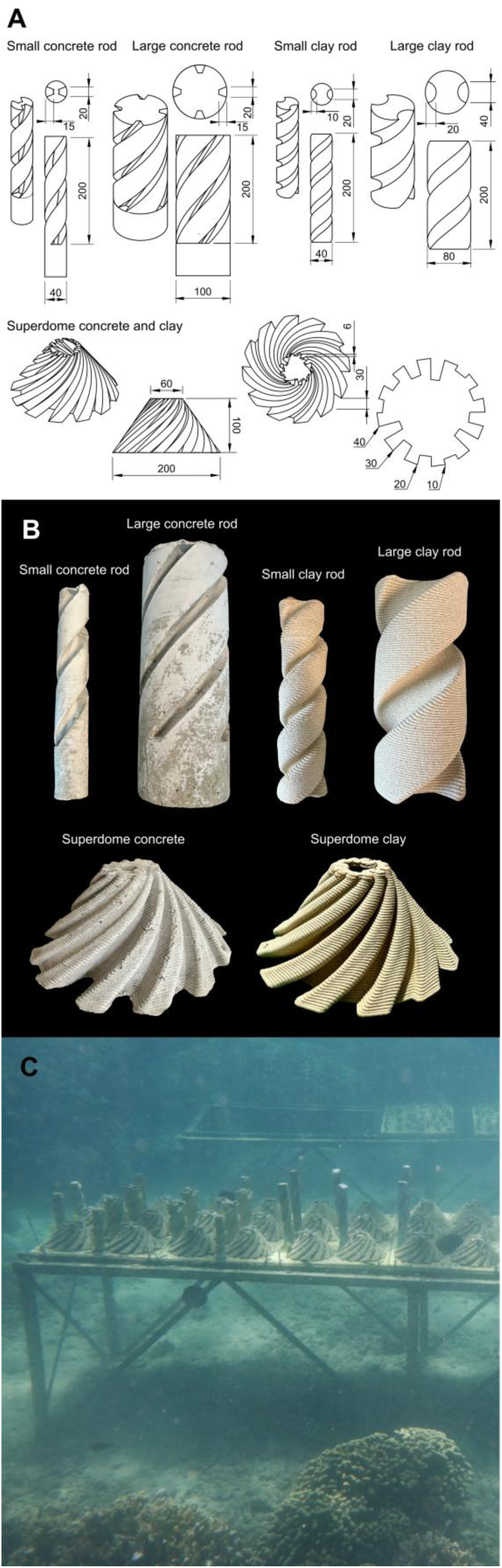
Designs of coral settlement modules and deployment. The designs demonstrate that helix recess design can be implemented across both dome and rod geometries, enabling comparative testing of settlement performance. (A) Dimensions of the six tested coral settlement modules (small concrete rod, large concrete rod, small clay rod, large clay rod, superdome concrete, and superdome clay) are presented in technical drawings (unit: mm). Designs were produced in concrete and clay. Rod designs all shared a height of 20 cm with varying diameters and recess dimensions. Superdomes had a conical shape with a base diameter of 20 cm, a height of 10 cm, and decreasing recess sizes towards the top. Recesses had four different depths decreasing towards the top with maximal depths of 1 cm, 2 cm, 3 cm, and 4 cm, respectively. Details of the technical dimensions are summarized in Table S2. (B) Photos of coral settlement modules in the six different designs tested. (C) Modules were deployed on tables at patch Reef 13, Kāneʻohe Bay at ∼2 m depth.

### 2.2 Module design and production

Six types of settlement modules were fabricated with two different module geometries (cylindrical rods and conical domes), different diameters (small and large rods), and two materials (cast in concrete and 3D-printed in clay), (Figure 1A, B). Details of the technical dimensions are provided in Table S1. Different rod diameters were tested to introduce controlled variation in macro-scale curvature, which affects near-surface flow separation, recirculation patterns, and shading. Rods with larger diameter generate broader low-velocity and light-exposed zones than rods with smaller diameter. It thus allows us to evaluate whether helix recess performance is robust across hydrodynamic and light environments. ‘Small rods’ had a base of 40 mm in diameter. ‘Large rods’ had a base of 80 mm (clay) or 100 mm (concrete) in diameter. All rods featured continuous spiral helix recesses (four recesses in the large concrete module, two recesses in all other modules) designed to mimic the successful ‘superdome’ recesses (Reichert et al., 2024). All cylindrical rods shared a height of 200 mm and featured a continuous spiral helix recess (Figure 1A). The conical modules were based on the ‘superdome’ modules used in previous studies (20 cm base diameter, 10 cm height, 12 helix recesses across four depths; Reichert et al., 2024) and were produced in both concrete (’Superdome concrete’) and clay (’Superdome clay’). The helix recesses on the rod modules were intended to mimic the successful settlement microhabitats identified in the superdomes (i.e., recesses with 17 mm depth and 15 mm width, Reichert et al., 2024).

The concrete settlement modules (‘small concrete rod’, ‘large concrete rod’, and ‘superdome concrete’) were fabricated through casting. For both concrete rod settlement models, the recess profiles were designed as trapezoidal shapes at ∼98° angle and 20 mm width and depth (Figure 1A). Settlement modules were designed in Autodesk Fusion 360 and master patterns were 3D printed in PLX material (PLA-derived bio-performance filament, BigRep GmbH, Germany) on a BigRep ONE.4 printer (BigRep GmbH), (Figure S1). Flexible molds were created from these master patterns using Mold Star™ 30 platinum-cure silicone rubber (Smooth-On, Inc., USA). The final concrete settlement modules were cast within these silicone molds using a high-strength concrete mix (Quikrete® Concrete Mix No. 1101). An attachment screw was embedded centrally during casting to facilitate underwater deployment. Following demolding, the cast settlement modules were cured for at least two days before deployment. The ‘superdome concrete’ settlement modules were fabricated in the same way, with the master pattern being a clay-printed and fired superdome.

The clay settlement modules (‘small clay’, ‘large clay’, and ‘superdome clay’) were fabricated using 3D printing. Designs from Autodesk Fusion 360 were exported as .stl files and processed using Simplify3D slicing software. Settlement modules were printed using a Delta Wasp 40100 3D printer (Wasp, Italy) with mid-fire pottery clay (Soldate 60, Aardvark Clay and Supplies, United States), prepared using a pugmill (Pugmill/Mixer NVS-07, Nidec, Japan). All clay settlement modules were printed at 110% scale to compensate for shrinkage during drying and firing. Clay was extruded through a 4 mm diameter nozzle, creating a distinct horizontal layer microstructure on surfaces (Figure S1, microstructure clay). The settlement modules were printed as hollow structures using the spiral vase mode, following the procedures developed in previous studies (Reichert et al., 2024). However, printing the target recess geometry on the cylindrical clay rods proved structurally unstable, so the recess profile for the clay rods was modified to a more rounded shape. Notably, the ‘large clay’ design was printed with wider recesses compared to the target dimensions. After printing, all clay settlement modules were air-dried for 7 to 10 days in a stable environment until bone-dry and were then fired in a kiln (Jupiter Sectional Kiln, L&L Kilns, United States) at ∼2030 °F (≈1100 °C) with a 2-hour hold (setting: cone 8).

### 2.3 Deployment and monitoring

Coral settlement modules were deployed on tables at patch Reef 13, in Kāneʻohe Bay, Oʻahu, Hawaiʻi (21.4504444° N, −157.7956111° W), at ∼2 m depth on 21 June 2024, one week before the anticipated mass spawning of *M. capitata* corals, to allow for biofilm conditioning on the structures prior to spawning. Settlement modules were deployed in an upright position onto experimental tables (Figure 1C). Coral recruitment on the modules was assessed at 1 week, 2 months, and 6 months post-spawning. For this, modules were transported back to the lab at Moku o Lo‘e, where recruits were counted via fluorescence detection (Sola Nightsea Blue Light, blue-light-blocking glasses). After counting, the modules were returned to the field site. The number of recruits was counted separately on outside surfaces and surfaces inside the recesses. Recruit counts were converted to recruit densities (per cm²) by standardizing to the location-specific surface area of each module type, determined in the 3D model designs. Further, to determine the planar recruit density, total numbers per module were standardized to the planar area of each module.

Recruitment on settlement modules was compared to recruitment on natural reef substrate. After two months, coral recruits were counted in 32 randomly selected 0.5 x 0.5 m quadrats in the nearby Barrier Reef in Kāneʻohe Bay (21.458853° N, 157.798046° W), using blue light fluorescence. The percent cover of benthic components (CCA, live coral, and rubble/sand) was visually estimated. The three-dimensional structure of each quadrat was reconstructed using photogrammetry from approximately 2500 overlapping images captured in a spiral pattern, following established procedures (Pizarro et al., 2017). Orthomosaics were built in Agisoft Metashape photogrammetric software (Roach et al., 2021). Recruitment density was then standardized to the available surface area by subtracting non-settlement areas (live coral, sand). Recruit density in the reef was similarly standardized to the three-dimensional surface area available, as well as to the planar surface area.

### 2.4 Statistical Analysis

Data were analyzed in R (v.4.5.0, R Core Team, 2025). Differences in recruit densities among the six structure types were assessed using pairwise Wilcoxon rank-sum tests with Holm correction. Comparisons between locations (’Inside’ vs. ‘Outside’) within each structure type were performed using Wilcoxon rank-sum tests. To assess how structural and material features influenced coral recruit density over time, generalized linear models were fitted separately for each timepoint using a Tweedie distribution with a log link function in the glmmTMB package (Brooks et al., 2017). The Tweedie family is well-suited for modeling continuous, non-negative data with a large proportion of zeros and a right-skewed distribution, such as coral recruit densities (recruits per cm²). Models included the fixed effects of shape (rod vs. superdome), recruit location (inside vs. outside), material (clay vs. concrete), diameter (large vs. small), and recess size (large vs. small). Model diagnostics were performed with the DHARMa package and included tests for overdispersion, zero inflation, and quantile deviations, none of which indicated violations of model assumptions (Hartig, 2025). Estimated marginal means (EMMs) and pairwise contrasts were calculated using the emmeans package (Lenth, 2025). Temporal changes in recruitment and post-settlement survival were quantified using recruit densities measured inside recesses. For each module type, recruit survival between consecutive time intervals was calculated as the percentage of recruit density retained. Survival rates among module types were compared using pairwise Wilcoxon rank-sum tests followed by Holm correction for multiple testing. In addition, a Tweedie generalized linear model with a log link function was fitted with recruit density as the response variable and timepoint, module type, and their interaction as fixed effects to test for time-by-module-type interactions. To compare artificial modules with natural reef surfaces, recruit densities on modules were contrasted with densities on reef quadrats standardized by three-dimensional habitat area and planar area derived from photogrammetry at the two-month time point. Wilcoxon rank sum tests with Holm correction were used to compare reef quadrats with settlement modules for both recess-based and planar densities.

## 3. Results

### 3.1 Overall recruitment dynamics

Coral recruitment occurred on all deployed structure types featuring helix recesses. While early settlement varied significantly among designs, long-term recruitment success converged across module types (Figure 2, Figure S2, Table S2). In particular initial settlement densities at 1 week post-spawning differed significantly between the structures (Wilcoxon rank-sum test, p<0.05). Recruit densities were highest on concrete structures, reaching densities up to 2.86 ± 0.77 recruits per cm² (inside the recesses) in the large concrete rod design (Figure 2A). Recruit densities declined markedly after 2 months due to post-settlement mortality. Recruit densities stabilized equally across all designs around 0.12 ± 0.04 recruits per cm² (Wilcoxon rank-sum test, p>0.05).

**Figure 2:**
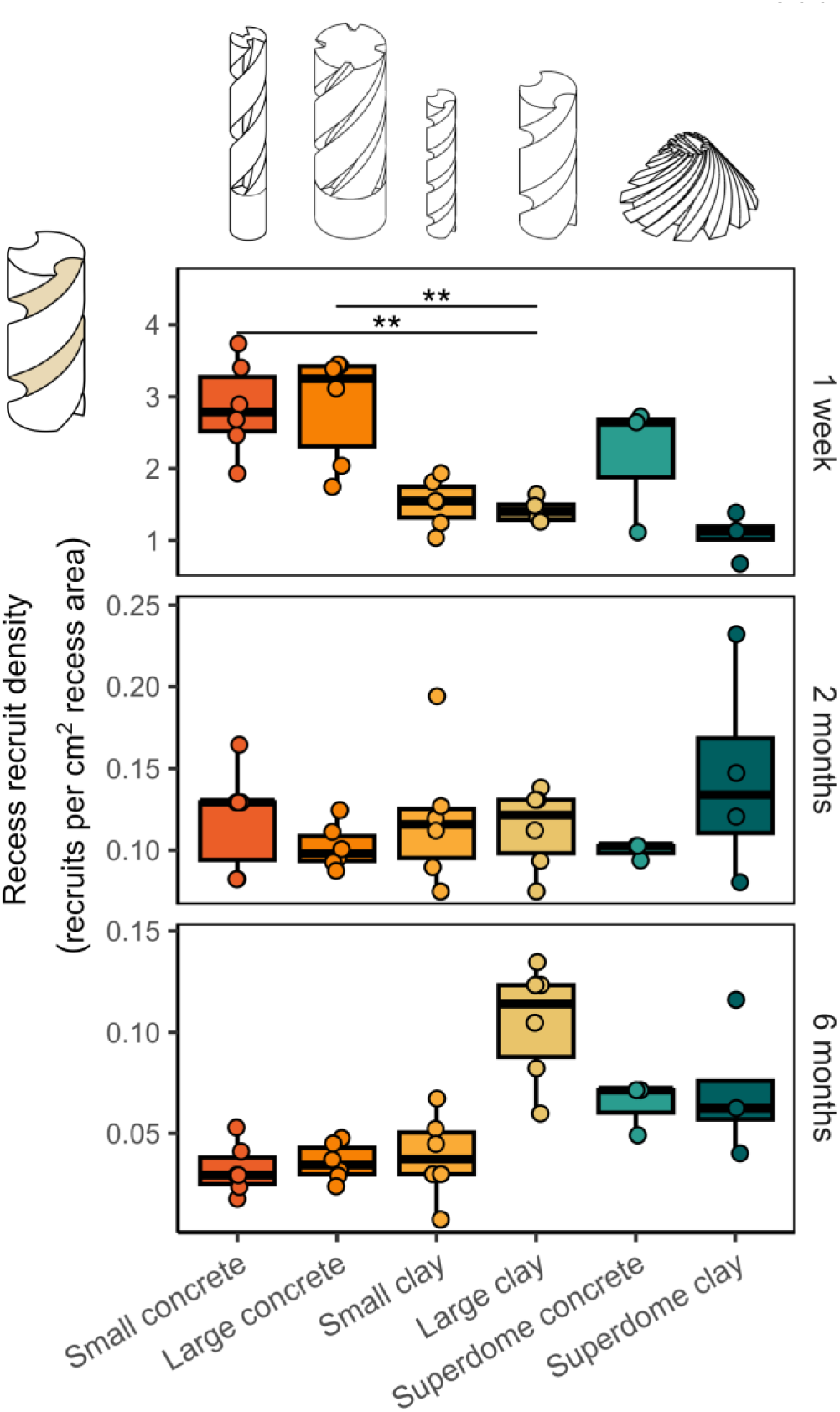
Coral recruitment on the six tested coral settlement modules (small concrete rod, large concrete rod, small clay rod, large clay rod, superdome concrete, and superdome clay) after 1 week, 2 months, and 6 months reveals that while early settlement varied significantly among designs, long-term recruitment success was comparable across all module types. Recruit densities are analyzed inside the recesses of the structure (recruits per cm^2^). Data are displayed as box-and-whisker plots with raw data points; lines indicate medians, boxes indicate the first and third quartile, and whiskers indicate ± 1.5 IQR. Asterisks indicate levels of statistical significance, derived from Wilcoxon rank-sum tests followed by Holm adjustment for multiple testing: * p ≤ 0.05, ** p ≤ 0.01, ***p ≤ 0.001.

Between months 2 and 6, densities showed slower declines on most structures, leading to values around 0.06 ± 0.03 recruits per cm² at the 6-month mark. Notably, recruit density rankings shifted compared to the initial settlement. In particular, the large clay design, which had the lowest initial recruitment, had the highest recruit densities after six months. On all settlement modules, recruit densities inside the recesses were considerably higher than on the total module, as only few recruits settled on the outside surfaces (Figure S2A, Table S3, Figure 4B). The relative trends among module shapes were consistent across these density metrics. Planar recruit densities revealed significantly higher densities on the rod-shaped designs compared to the superdome designs, with highest densities found on the small concrete rod design (Figure S2C).

Tweedie generalized linear models revealed factors that influence coral settlement and survival (Table 1). The recruitment location within the module (inside recess vs. outside of recess surfaces) was by far the most consistent and influential factor across all time points. Recruits were significantly more abundant on the inside recess surfaces than on the exposed outside surfaces at 1 week, 2 months, and 6 months (all p < 0.0001). The effect was strongest at one week, indicating a strong initial preference for the protected microhabitats inside recesses. Recruit densities after one week were higher on rods than on superdomes, showing an early effect of module shape (p = 0.0152). However, the effect weakened over time and was not visible after 2 or 6 months. Material affected early settlement, with concrete supporting higher recruit densities than clay after 1 week (p < 0.0001). However, this difference disappeared at 2 months and 6 months, suggesting that while concrete may promote initial settlement, clay supports comparable long-term outcomes. Recess size had no detectable effect at month 1 or 2 but became a significant factor after 6 months. Larger recesses supported higher recruit densities (p = 0.0013), indicating a delayed benefit of larger microhabitats for post-settlement survival and persistence. The rod diameter had no significant effect at any time point. Although rods with larger diameter often showed slightly higher densities, these differences were small and not significant, indicating that module diameter played only a minor role.

**Table 1:**
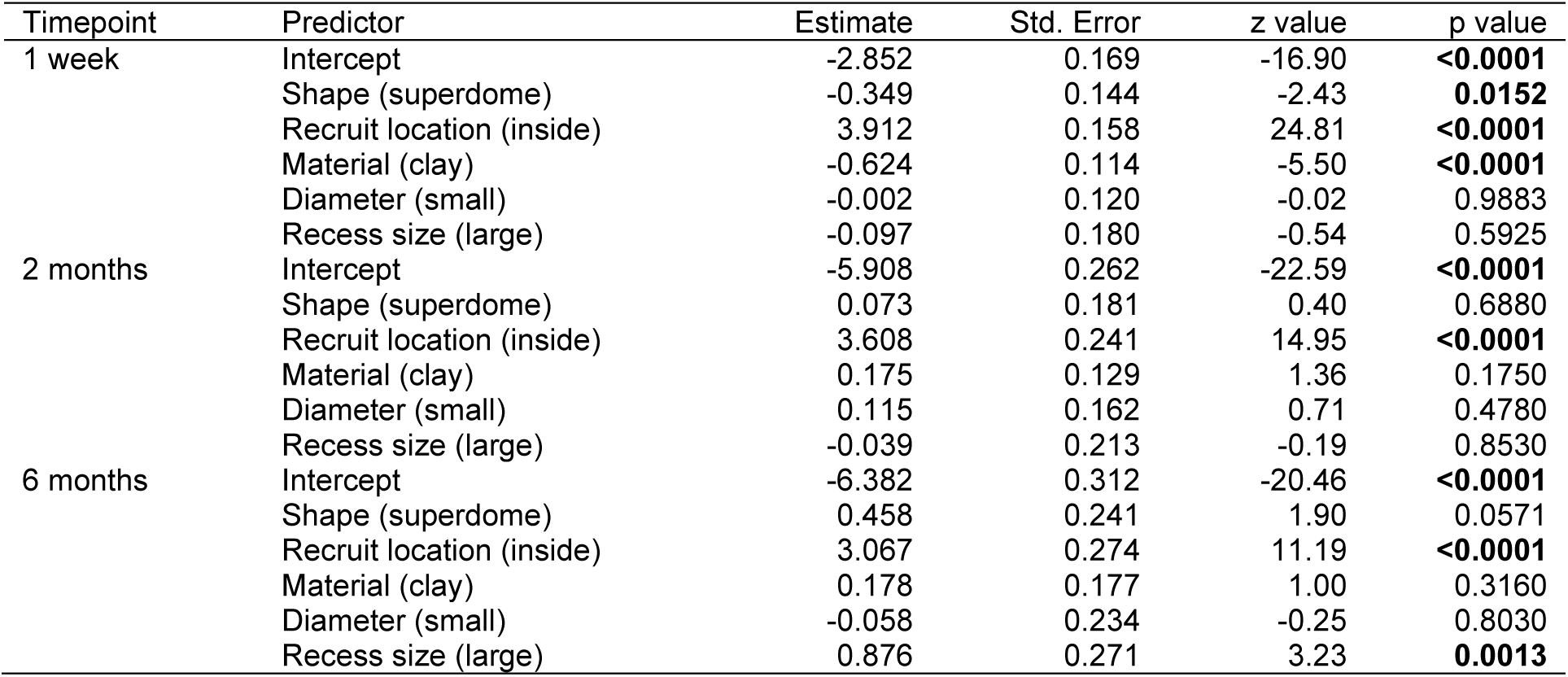
Statistical results of Tweedie generalized linear models assessing the effects of shape, recruit location, material, diameter, and recess size on coral recruit density (recruits per cm²) after 1 week, 2 months, and 6 months. Model-estimated coefficients (log scale), standard errors, z-values, and p-values are given for each predictor. Significant effects (p < 0.05) are highlighted in bold.

### 3.2 Post-settlement survival

Despite slight differences in the recess geometry between concrete rods (designed to optimal dimensions) and clay rods (rounded recesses constrained by 3D clay printing), the convergence of recruit densities by 2–6 months suggests that the overall helix recess design is robust and most designs showed similar survival rates (Figure 3). Only the large clay rod design stood out. While having wider recesses, this specific design showed minimal decline in recruit density between the 2-month and 6-month time points, indicating enhanced recruit survival during this phase compared to other designs (Figure 3B, Wilcoxon rank-sum test, p=0.0320, Table S4). A Tweedie interaction model including a timepoint x module type interaction confirmed the same overall pattern, with strong effects of timepoint (χ²₂ = 3244.14, p < 0.001), module type (χ²₅ = 16.51, p = 0.0055), and their interaction (χ²₁₀ = 130.23, p < 0.001), (Table S5).

**Figure 3:**
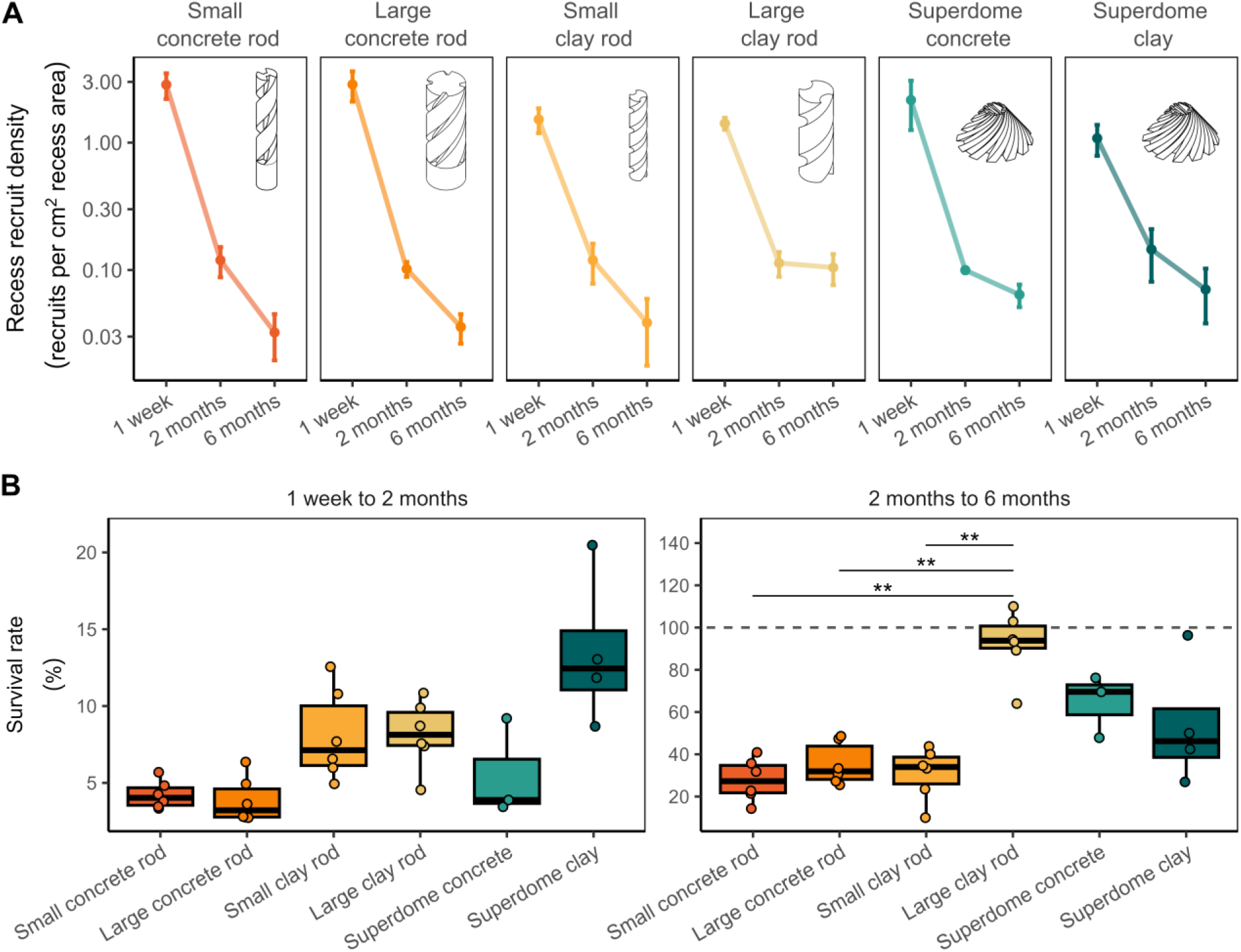
Coral recruit survival on different settlement modules. Survival rates suggest that the overall helix design concept is robust and most designs showed similar survival rates. (A) Declines in recruit densities (recruits per cm^2^ inside the recesses) on different designs in concrete and clay (small concrete rod, large concrete rod, small clay rod, large clay rod, superdome concrete, and superdome clay) from 1 week to 2 months, and to 6 months. Data are displayed as line plots with mean ± standard deviation. (B) Survival rate of recruits on the six different shapes from 1 week to 2 months, and from 2 months to 6 months. Data are displayed as box-and-whisker plots with raw data points; lines indicate medians, boxes indicate the first and third quartile, and whiskers indicate ± 1.5 IQR. Asterisks indicate levels of statistical significance, derived from Wilcoxon rank-sum tests followed by Holm adjustment for multiple testing: * p ≤ 0.05, ** p ≤ 0.01, ***p ≤ 0.001. The dashed line marks 100% survival.

#### Comparison to natural reef recruitment

To assess the performance of the tested coral settlement structures in comparison to natural reef conditions, recruit densities after two months were compared to recruitment on natural substrate at a nearby reef site (assessed through recruit counts in 32 quadrants of 0.5 m x 0.5 m). Mean recruit density on the reef was 25 ± 24 recruits per m² relative to its 3D surface and 30 ± 28 recruits per m² planar area, calculated based on available benthic settlement surface area (Figure 4A). In contrast, recruit densities on the modules were significantly higher (Wilcoxon rank-sum tests, p < 0.001, Holm-corrected; Table S6). Recruit densities inside the helix recesses of all tested module types were ∼50-fold higher at 1163 ± 345 recruits per m². Recruit densities on planar areas were ∼300-fold higher, with 8269 ± 6443 recruits per m² observed.

**Figure 4:**
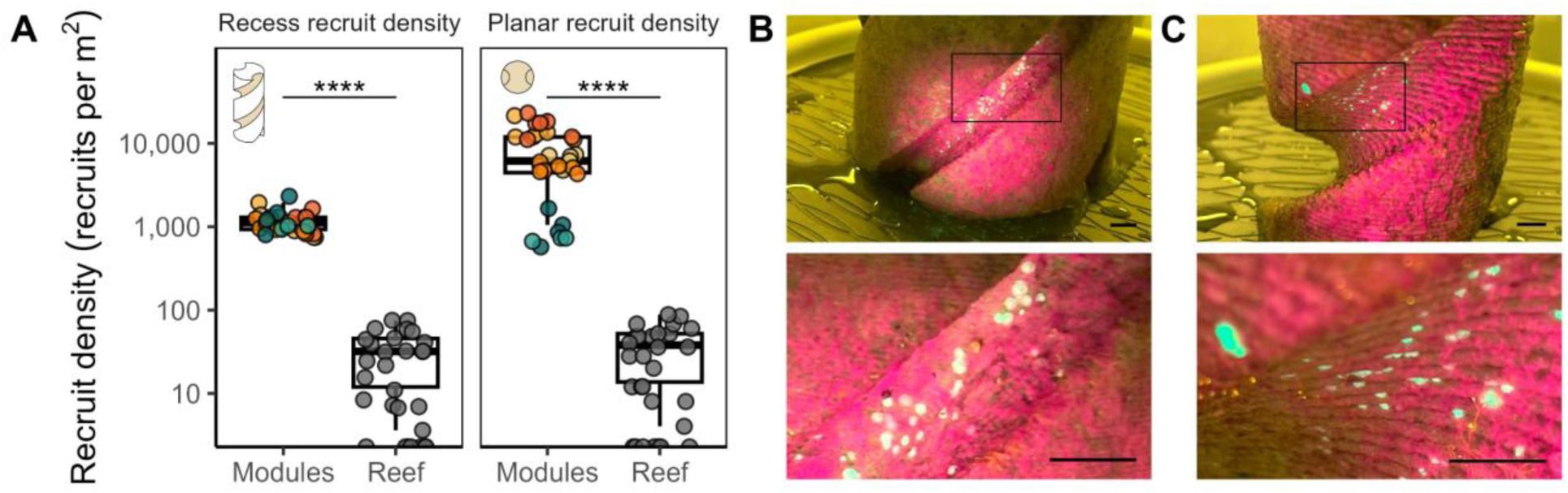
Coral recruitment on coral settlement modules was higher than recruitment on natural reef substrate in an adjacent reef after 2 months, assessed through recruit counts in 32 quadrants of 0.5 m x 0.5 m. (A) Recruit densities (recruits per m^2^) in the recesses and the 3D reef structure derived from photogrammetry and planar recruit densities are compared. Data are displayed as box-and-whisker plots with raw data points; lines indicate medians, boxes indicate the first and third quartiles, and whiskers indicate ± 1.5 IQR. Asterisks indicate levels of statistical significance, derived from Wilcoxon rank-sum tests: * p ≤ 0.05, ** p ≤ 0.01, ***p ≤ 0.001. (B) Photos of 1-week-old recruits inside the recesses of the large concrete design. Scale bars represent 1 cm. Recruits are green-fluorescent, excited via blue light, and visualized through yellow filters. (C) Photos of 1-week-old recruits inside the recesses of the large clay design, respectively.

## 4. Discussion

This study evaluated the transfer of optimized helix recess designs from conical domes to pylon-like, cylindrical rods and examined the influence of module geometry and material on coral recruitment. Our findings confirm the effectiveness of the helix design on cylindrical structures, demonstrate the robustness of the design across different materials, and provide new insights into how recess dimensions affect post-settlement survival.

### 4.1 Successful design transfer to cylinders

Similar recruitment densities on cylindrical rods and conical superdomes, especially after initial mortality, show that the helix recess design transfers successfully between module geometries. The helix recesses promote larval entrainment by creating small recirculation zones and reducing flow inside recesses, conditions that favor settlement (Hata et al., 2017; Levenstein et al., 2022a). Our results show that these processes work equally well in the recesses of the cylindrical rods and conical domes tested here. The recess dimensions of 1–10 mm match those preferred by coral larvae in other experiments (Doropoulos et al., 2016; Rahnke et al., 2022; Ramsby et al., 2024) and on natural reefs (Brandl et al., 2014; Edmunds et al., 2004). Recruits preferred inner recess surfaces over outer exposed areas, underlining that these features provide crucial settlement microhabitats on cylindrical structures.

### 4.2 Design robustness across materials

The helix recess design was the main driver of recruitment success. Material influenced initial settlement, with slightly higher densities on concrete, but this difference disappeared over time. The used concrete and clay modules differed in several physical properties relevant to early benthic colonization. Concrete surfaces exhibited higher microscale roughness but lower porosity, while 3D-printed and fired clay produced smoother, ridged microtopography with higher microporosity. Clay is chemically inert after firing, whereas concrete retains calcium-rich mineral phases that can influence microbial films and settlement cues. The two materials also differed in color and optical properties. Concrete modules were medium to dark grey, generating lower reflectance and darker recess interiors, while the light-beige clay surfaces produced higher reflectance and brighter, more diffuse light within recesses. We did not test material chemistry directly, such as potential leaching or pH modification by concrete, and our interpretations therefore rely on qualitative physical properties observed. However, such chemical effects could influence microbial film development and settlement cues and represent an important avenue for future work.

Coral larvae are known to settle on a wide range of materials, including coral skeletons, terracotta, cement, glass, and PVC (Babcock and Mundy, 1996; Hunte and Wittenberg, 1992; Soong et al., 2003; Wilson and Harrison, 1998). While some studies report preferences for ceramic tiles or coral skeletons, site-specific environmental conditions usually have a stronger influence on recruitment patterns (Burt et al., 2009). Initial settlement was slightly higher on concrete modules, a pattern consistent with material-driven differences documented in previous studies (Babcock and Mundy, 1996; Burt et al., 2009). Small deviations in recess geometry between concrete and clay rods influence near-surface hydrodynamics which can alter larval contact rates (Hata et al., 2017; Levenstein et al., 2022a). Surface chemistry and microscale porosity also vary between concrete and clay, and such differences are known to affect settlement cues, biofilm formation, and juvenile survival (Hoog Antink et al., 2018; Natanzi et al., 2021; Petersen et al., 2005). In addition, smoother or less porous materials can reduce long-term attachment strength, making recruits more susceptible to dislodgement (Carl et al., 2012). These factors provide a mechanistic basis for the initially higher settlement on concrete and the subsequent convergence of recruit densities across materials. In summary, these findings demonstrate that helix recess designs robustly enhance coral settlement and survival across different materials, such as clay and concrete. Concrete offers advantages in terms of cost, durability, and scalability (Spieler et al., 2001; Knoester et al., 2024). Since no differences in long-term survival were observed, concrete appears to be a suitable material for producing coral settlement modules for reef restoration.

### 4.3 Trade-offs between settlement and survival based on recess size

The large clay rods had wider, rounded recesses because of printing limitations. Despite lower initial settlement, they maintained the highest recruit densities between two and six months. This suggests that larger recesses can enhance post-settlement survival. This result complements our earlier finding that narrow recesses promote early settlement (Reichert et al., 2025), but indicates that wider recesses improve persistence under field conditions. Coral larvae are strongly influenced by microscale hydrodynamics and substrate geometry: Small recesses generate recirculation zones that entrain larvae that have very limited swimming ability (Hata et al., 2017; Levenstein et al., 2022a). These sheltered spaces also reduce early exposure to grazers (Brandl et al., 2014; Doropoulos et al., 2016; Nozawa, 2008) and sediment movement (Babcock and Davies, 1991; Hodgson, 1990), both important sources of early mortality. In contrast, wider recesses provide more space, higher light penetration, and improved water exchange, conditions known to support juvenile coral respiration, feeding, and skeletal growth (Brunner et al., 2022; Mass et al., 2010; Sebens et al., 1997). Larger recesses also buffer recruits from competitive overgrowth by turf algae or colonial invertebrates by increasing physical separation (Brandl et al., 2014; Chadwick and Morrow, 2011; Doropoulos et al., 2016). Together, these mechanisms explain why small recesses are advantageous for settlement, whereas larger recesses promote persistence across later developmental stages. In addition to these benefits, wider recesses may also introduce trade-offs. Larger recesses may therefore experience reduced internal flushing, which can promote sediment retention and microbial shifts. Such low-flow pockets consistently accumulate fine sediments, which impairs larval settlement and post-settlement survival (Hodgson, 1990; Ricardo et al., 2017; Weber et al., 2012). While such effects were not evident within the six-month timeframe of our experiment, they may become relevant under higher sedimentation regimes or longer deployments. Considering these potential constraints will be important when optimizing recess dimensions for both early settlement and longer-term module performance. Therefore, future work should test a wider range of recess dimensions to identify designs that balance settlement and survival.

### 4.4 Recess functionality and module size effects

Coral larvae were consistently found inside the recesses, confirming that helix recesses function as key settlement microhabitats, including on cylindrical substrates. This preference was strongest after one week, indicating an early selection for protected areas. Recruitment did not differ between small and large rod diameters, indicating that the tested centimeter-scale differences in curvature did not alter settlement outcomes. This aligns with studies showing that coral larval responses are primarily driven by microtopography rather than macro-scale substrate curvature (Doropoulos et al., 2016; Levenstein et al., 2022a; Petersen et al., 2005). The reduced settlement and survival on the outside surfaces are likely linked to the increased grazing pressure and competition from benthic algae on these surfaces (Brandl et al., 2014; Doropoulos et al., 2016). These results underline the functional value of helix recesses as a protective habitat for coral recruits.

### 4.5 Implications for coastal engineering and reef restoration

This study demonstrates that helix recesses enhance coral recruitment on cylindrical structures. The substantially higher settlement observed on the helix modules compared with nearby natural reef surfaces underscores their potential value for restoration. Settlement densities on natural reefs primarily reflect the availability of suitable settlement microhabitats (Vermeij and Sandin, 2008; Doropoulos et al., 2017). Although negative density dependence occurs during early post-settlement stages (Bramanti and Edmunds, 2016), high initial settlement is still necessary for populations to persist, because most juveniles die during the earliest weeks. The helix recesses create a continuous high-density of preferred microhabitats that promote larval entrainment and retention. Therefore, the resulting 300-fold differences in settlement density reflect the success of the structures and demonstrate that microhabitat-focused design can reliably amplify larval settlement. Consequently, integrating such features into coastal and restoration infrastructure can meaningfully accelerate reef recovery.

The design was effective across both concrete and clay, allowing material choice to depend on durability, cost, or environmental footprint. Helix recesses can therefore be integrated into cylindrical and pylon-like elements, such as pilings, tetrapods, seawalls, or groin foundations, commonly used in coastal engineering and shore protection. With that, coastal structures could be adapted to provide ecological function as coral settlement habitat and serve as part of nature-based solutions and hybrid infrastructure, where shoreline protection and reef restoration are achieved simultaneously. This dual function improves the cost-effectiveness of restoration by building on established engineering practice. Over longer periods, this method could also improve the stability and lifespan of coastal infrastructure. Early successional stages may enhance functionality by stabilizing surfaces and promoting biological binding (Rinkevich, 2021), as corals can grow out of the recesses and begin to encrust exposed surfaces within months (Reichert et al., 2025, Schiettekatte et al., in prep). Mature settlement communities can modify hydrodynamics, sediment retention, and surface accessibility. With larval settlement and continuous growth of previously settled reef-building corals, these settlement modules (1) demonstrate self-regenerating properties as they can mitigate and repair damage caused e.g. through storms or anchor damage. (2) Rising sea levels can potentially reduce the effectiveness of coastal protection infrastructure (Griggs & Reguero, 2021; Storlazzi et al., 2021; Toth et al., 2023) by increasing the water height above the structure and affecting other relevant hydrodynamic parameters (Quataert et al., 2015). Compared to solely “gray” coastal protection solutions, microhabitats facilitate continuous vertical coral growth and reef accretion and can thus provide a natural long-term solution to dynamically adjust to a rising sea level. Recesses also gradually fill with other benthic organisms, such as calcareous encrusters, sponges, and bryozoans (observational data), which can reduce available settlement habitat but also generate additional microtopography that supports repeated settlement events. Further, accumulated sediments might reduce the available space for successive settlement (Hodgson, 1990; Ricardo et al., 2017; Weber et al., 2012). Assessing these long-term dynamics under high-energy and turbid coastal conditions will be essential for optimizing recess geometry for real-world applications.

An important next step is to evaluate how recesses perform over extended periods, including whether they can support repeated settlement events that build self-sustaining coral populations. Furthermore, this study emphasizes that recess design must strike a balance between features that support initial settlement and those that promote survival. Varying recess size across modules may improve performance across coral life stages. Future research should therefore systematically investigate larger recess geometries, as well as a broader range of coral species and reef environments, to establish general design rules for restoration-oriented engineering.

In summary, cylindrical elements with helix recesses are scalable and low-cost. They can be mass-produced in concrete and incorporated into coastal engineering projects to combine shoreline protection with reef restoration.

## Acknowledgements

We thank Guan-Yan Chen, Marion Chapeau, and Corryn Haynes from the Geometric Ecology Lab for their support during the fieldwork. We thank the students of the 2024 MBIO640 class at the University of Hawaiʻi for their help in collecting the reef recruitment data. This work was funded by the Defense Advanced Research Projects Agency (Contract No. HR001122C0134), the National Science Foundation (1948946), and the HIMB Director’s Innovation Fund.

## Author contributions

Conceptualization: JR, HJ, and JSM

Methodology: JR, HJ, and JSM

Data curation: JR, ADN, LHR

Formal analysis: JR

Investigation: JR, HJ, MER, ADN, LHR, and CD

Resources: JSM, CD

Visualization: JR

Writing—original draft: JR

Writing—review & editing: JR, MER, JSM

## Data and materials availability

Data and scripts will be made available from GitHub.

## Declaration of generative AI and AI-assisted technologies in the manuscript preparation process

During the preparation of this work, Gemini (Google AI) and ChatGPT (OpenAI) were used to improve the clarity and language of the manuscript. The authors thoroughly reviewed all content and take full responsibility for the published article.

