## Supplementary material for "Cylindrical, pylon-like structures with helix recesses enhance coral larval recruitment": Figure S1

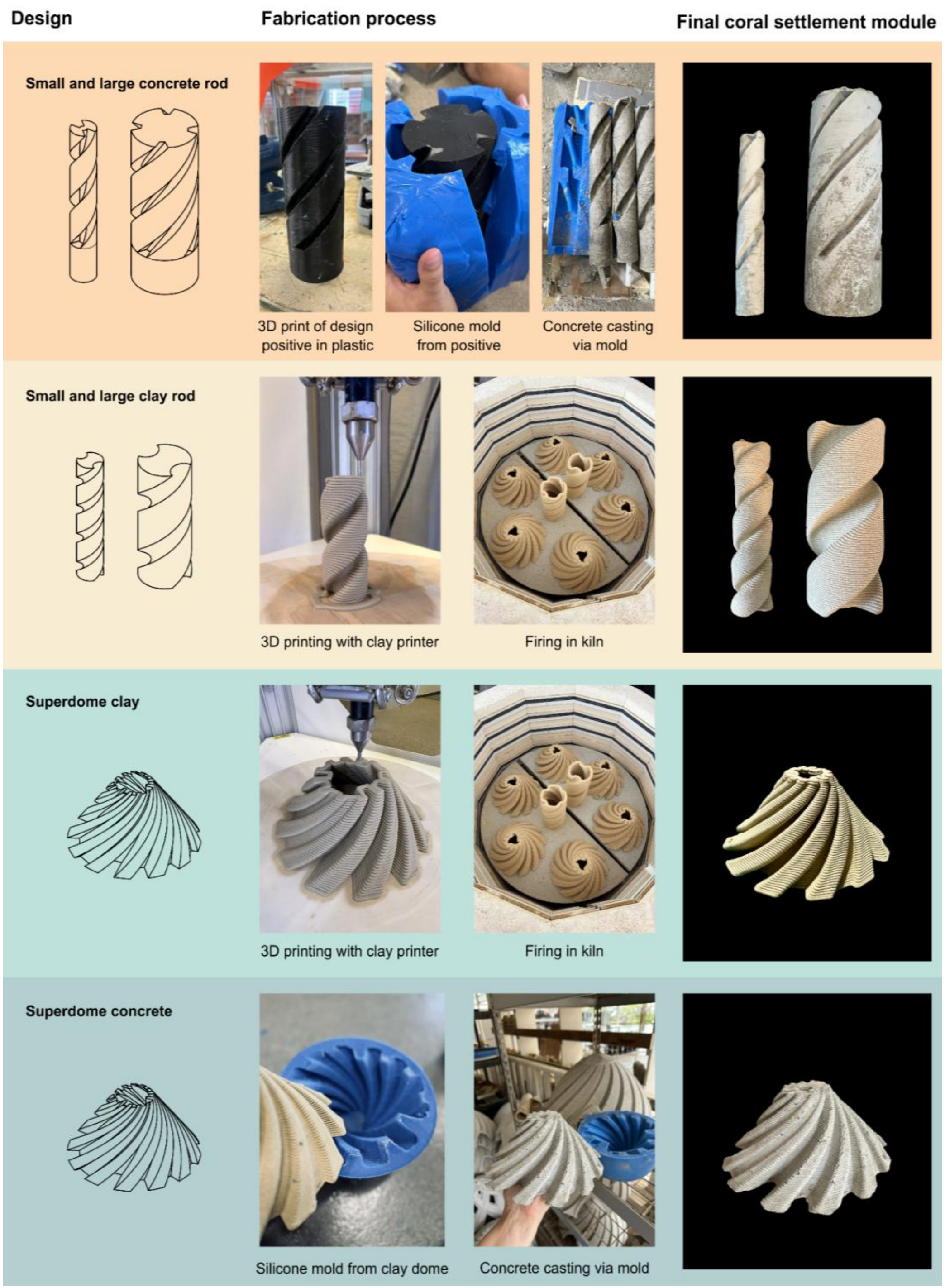


**Figure S1: Different fabrication processes of the tested designs.** Small and large concrete rods were fabricated by 3D printing plastic master patterns from which silicone molds were created. These molds were used to cast the final settlement modules in concrete. Small and large clay rods were directly 3D printed in clay, dried, and fired in a kiln. Superdome clay modules were created accordingly. Superdome concrete modules were fabricated by creating a silicone mold from a fired clay dome. The mold was used to cast the final settlement modules in concrete.

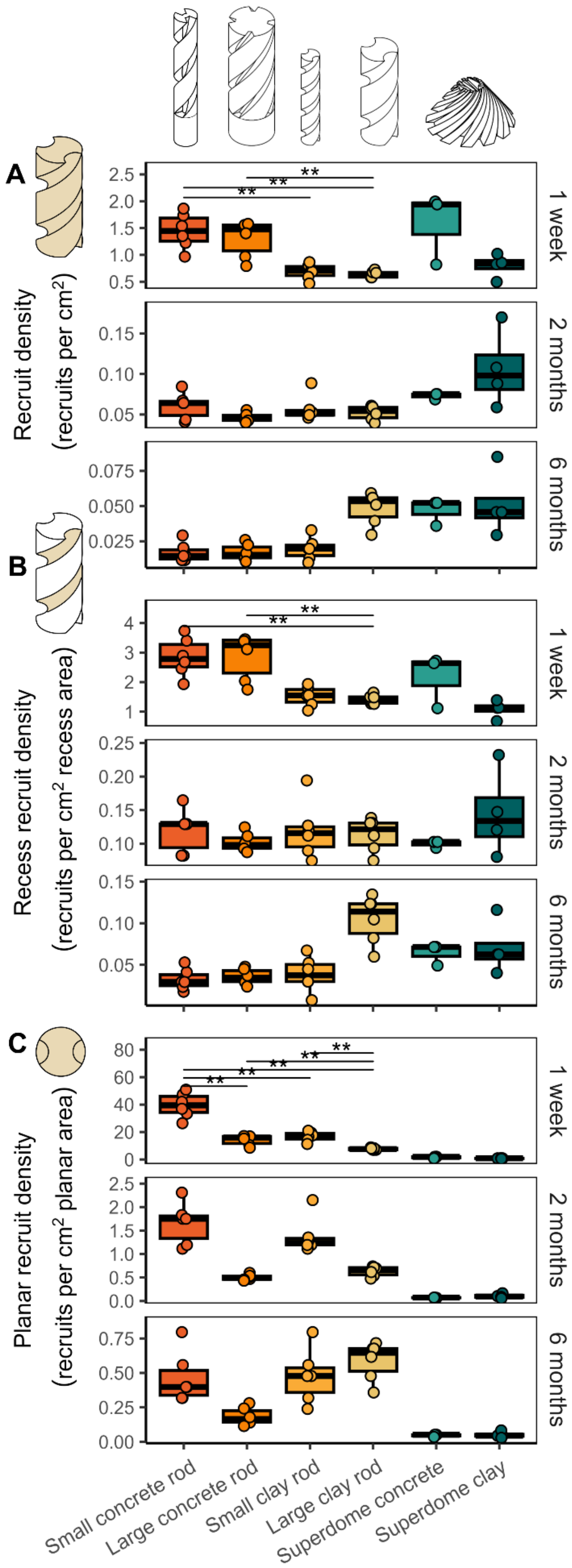


**Figure S2**: Coral recruitment on the six tested coral settlement modules (small concrete rod, large concrete rod, small clay rod, large clay rod, superdome concrete, and superdome clay) after 1 week, 2 months, and 6 months reveals that while early settlement varied significantly among designs, long-term recruitment success was comparable across all module types. (A) Recruit densities (recruits per $cm^2$) on the full structure. (B) Recruit densities (recruits per $cm^2$) inside the recesses of the structure. (C) Recruit densities (recruits per $cm^2$) per planar area of the structure. Data are displayed as box-and-whisker plots with raw data points; lines indicate medians, boxes indicate the first and third quartile, and whiskers indicate ± 1.5 IQR. Asterisks indicate levels of statistical significance, derived from Wilcoxon rank-sum tests followed by Holm adjustment for multiple testing: * $p \leq 0.05$, ** $p \leq 0.01$, ***$p \leq 0.001$.

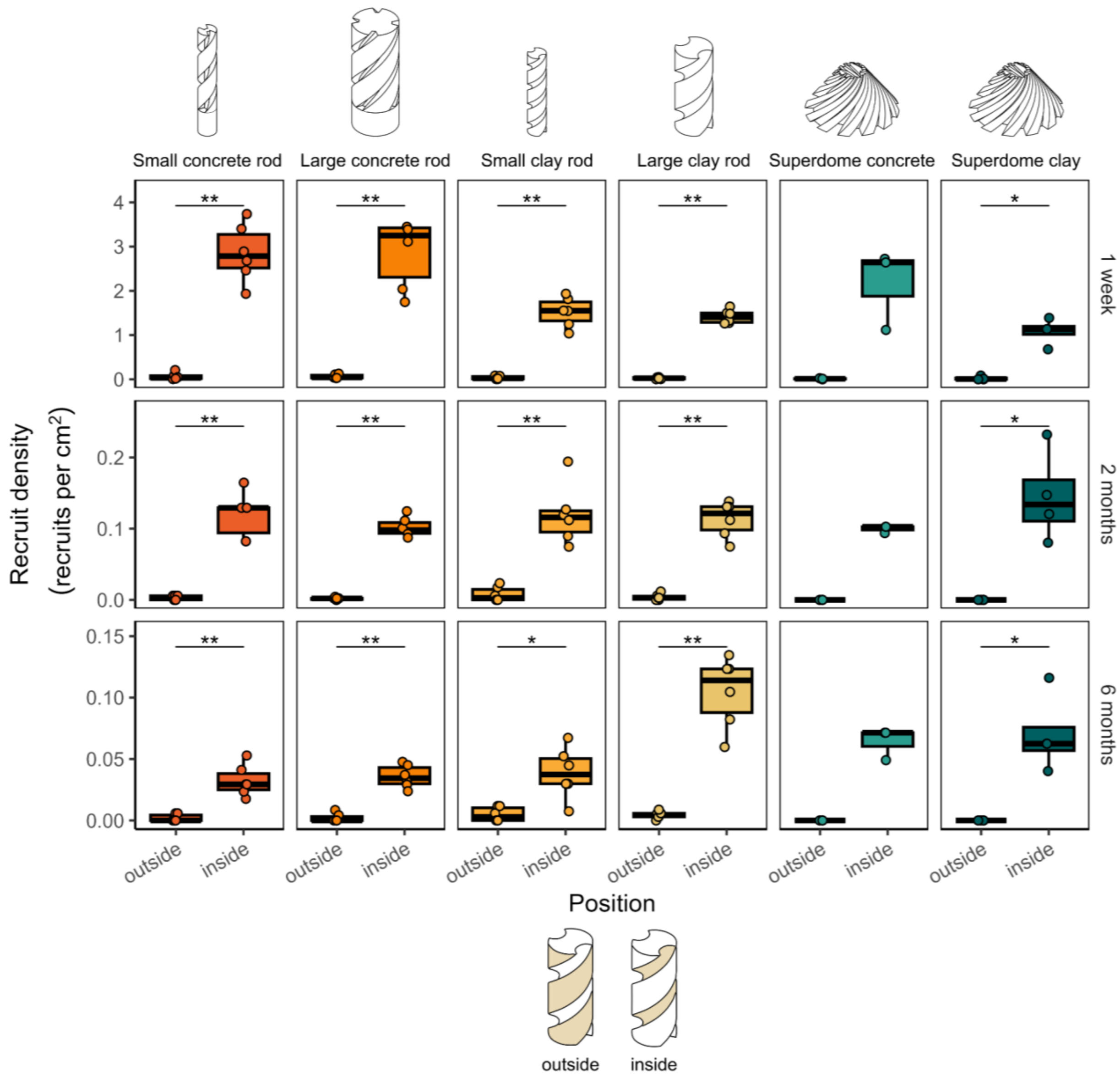


**Figure S3: Differences in coral recruitment on inside and outside surfaces.** Recruit densities (recruits per $cm^2$) of the six tested coral settlement modules (small concrete rod, large concrete rod, small clay rod, large clay rod, superdome concrete, and superdome clay) are standardized to the respective surface areas. Data are displayed as box-and-whisker plots with raw data points; lines indicate medians, boxes indicate the first and third quartile, and whiskers indicate ± 1.5 IQR. Asterisks indicate levels of statistical significance, derived from Wilcoxon rank-sum tests: * p ≤ 0.05, ** p ≤ 0.01, ***p ≤ 0.001.

**Table S1:** Dimensional parameters of the six settlement module designs used in this study. All cylindrical modules contain uniform recess geometry along their entire height, whereas the conical superdome designs incorporate four helix recess-depth levels (10, 20, 30, and 40 mm) that decrease toward the top.

| Module type | Geometry | Material | Diameter (mm) | Height (mm) | Recesses (numbers) | Recess depth (mm) | Recess width (mm) |
|---|---|---|---|---|---|---|---|
| Small concrete rod | Cylinder | Concrete | 40 | 200 | 2 | 15 | 20 |
| Large concrete rod | Cylinder | Concrete | 100 | 200 | 4 | 20 | 20 |
| Small clay rod | Cylinder | Clay | 40 | 200 | 2 | 10 | 20 |
| Large clay rod | Cylinder | Clay | 80 | 200 | 2 | 20 | 40 |
| Superdome concrete | Conical dome | Concrete | 200 | 100 | 12 | 10, 20, 30, 40 | 30–6 |
| Superdome clay | Conical dome | Clay | 200 | 100 | 12 | 10–2, 20–5, 30–8, 40–11 | 30–6 |

**Table S2**: Differences in recruit densities (number of recruits per $cm^2$) per module (standardized to the total surface area of the structure), recess recruit density (standardized to the surface area of the inside recesses of the structure), and planar recruit densities (standardized to the planar surface area of each module) between the six different shapes (superdome clay, superdome concrete, large clay rod, small clay rod, large concrete rod, small concrete rod) after one week, two months, and six months. Results are derived from Wilcoxon rank-sum tests followed by Holm adjustment for multiple testing. Number of replicates per test group (n1 and n2), test statistic (*W*), and p-value are given with bold values indicating significance ($p < 0.05$).

| Parameter | Timepoint | Contrast | | | n1 | n2 | Statistic | p |
|---|---|---|---|---|---|---|---|---|
| Recruit density per module | 1 week | Small concrete rod | – | Large concrete rod | 6 | 6 | 21 | 1 |
| | | Small concrete rod | – | Small clay rod | 6 | 6 | 36 | **0.0320** |
| | | Small concrete rod | – | Large clay rod | 6 | 6 | 36 | **0.0320** |
| | | Small concrete rod | – | Superdome concrete | 6 | 3 | 6 | 1 |
| | | Small concrete rod | – | Superdome clay | 6 | 4 | 23 | 0.2090 |
| | | Large concrete rod | – | Small clay rod | 6 | 6 | 34 | 0.1040 |
| | | Large concrete rod | – | Large clay rod | 6 | 6 | 36 | **0.0320** |
| | | Large concrete rod | – | Superdome concrete | 6 | 3 | 5 | 1 |
| | | Large concrete rod | – | Superdome clay | 6 | 4 | 20 | 0.9120 |
| | | Small clay rod | – | Large clay rod | 6 | 6 | 25 | 1 |
| | | Small clay rod | – | Superdome concrete | 6 | 3 | 1 | 0.4280 |
| | | Small clay rod | – | Superdome clay | 6 | 4 | 7 | 1 |
| | | Large clay rod | – | Superdome concrete | 6 | 3 | 0 | 0.2380 |
| | | Large clay rod | – | Superdome clay | 6 | 4 | 6 | 1 |
| | | Superdome concrete | – | Superdome clay | 3 | 4 | 9 | 1 |
| | 2 months | Small concrete rod | – | Large concrete rod | 6 | 6 | 28 | 1 |
| | | Small concrete rod | – | Small clay rod | 6 | 6 | 20 | 1 |
| | | Small concrete rod | – | Large clay rod | 6 | 6 | 26 | 1 |
| | | Small concrete rod | – | Superdome concrete | 6 | 3 | 3 | 1 |
| | | Small concrete rod | – | Superdome clay | 6 | 4 | 4 | 1 |
| | | Large concrete rod | – | Small clay rod | 6 | 6 | 8 | 1 |
| | | Large concrete rod | – | Large clay rod | 6 | 6 | 10 | 1 |
| | | Large concrete rod | – | Superdome concrete | 6 | 3 | 0 | 0.3850 |
| | | Large concrete rod | – | Superdome clay | 6 | 4 | 0 | 0.1430 |
| | | Small clay rod | – | Large clay rod | 6 | 6 | 18 | 1 |
| | | Small clay rod | – | Superdome concrete | 6 | 3 | 3 | 1 |
| | | Small clay rod | – | Superdome clay | 6 | 4 | 2 | 0.4640 |
| | | Large clay rod | – | Superdome concrete | 6 | 3 | 0 | 0.3850 |
| | | Large clay rod | – | Superdome clay | 6 | 4 | 2 | 0.4570 |
| | | Superdome concrete | – | Superdome clay | 3 | 4 | 3 | 1 |
| | 6 months | Small concrete rod | – | Large concrete rod | 6 | 6 | 18 | 1 |
| | | Small concrete rod | – | Small clay rod | 6 | 6 | 15 | 1 |
| | | Small concrete rod | – | Large clay rod | 6 | 6 | 0 | 0.0730 |
| | | Small concrete rod | – | Superdome concrete | 6 | 3 | 0 | 0.2430 |
| | | Small concrete rod | – | Superdome clay | 6 | 4 | 0 | 0.1600 |

| | | | | | | | | |
|---|---|---|---|---|---|---|---|---|
| | | Large concrete rod | – | Small clay rod | 6 | 6 | 15 | 1 |
| | | Large concrete rod | – | Large clay rod | 6 | 6 | 0 | 0.0730 |
| | | Large concrete rod | – | Superdome concrete | 6 | 3 | 0 | 0.2430 |
| | | Large concrete rod | – | Superdome clay | 6 | 4 | 0 | 0.1600 |
| | | Small clay rod | – | Large clay rod | 6 | 6 | 1 | 0.1040 |
| | | Small clay rod | – | Superdome concrete | 6 | 3 | 0 | 0.2430 |
| | | Small clay rod | – | Superdome clay | 6 | 4 | 1 | 0.2430 |
| | | Large clay rod | – | Superdome concrete | 6 | 3 | 11 | 1 |
| | | Large clay rod | – | Superdome clay | 6 | 4 | 14 | 1 |
| | | Superdome concrete | – | Superdome clay | 3 | 4 | 7 | 1 |
| Recess recruit density | 1 week | Small concrete rod | – | Large concrete rod | 6 | 6 | 17 | 1 |
| | | Small concrete rod | – | Small clay rod | 6 | 6 | 35 | 0.0560 |
| | | Small concrete rod | – | Large clay rod | 6 | 6 | 36 | **0.0320** |
| | | Small concrete rod | – | Superdome concrete | 6 | 3 | 13 | 1 |
| | | Small concrete rod | – | Superdome clay | 6 | 4 | 24 | 0.1050 |
| | | Large concrete rod | – | Small clay rod | 6 | 6 | 34 | 0.1040 |
| | | Large concrete rod | – | Large clay rod | 6 | 6 | 36 | **0.0320** |
| | | Large concrete rod | – | Superdome concrete | 6 | 3 | 14 | 1 |
| | | Large concrete rod | – | Superdome clay | 6 | 4 | 24 | 0.1050 |
| | | Small clay rod | – | Large clay rod | 6 | 6 | 22 | 1 |
| | | Small clay rod | – | Superdome concrete | 6 | 3 | 5 | 1 |
| | | Small clay rod | – | Superdome clay | 6 | 4 | 20 | 0.9120 |
| | | Large clay rod | – | Superdome concrete | 6 | 3 | 6 | 1 |
| | | Large clay rod | – | Superdome clay | 6 | 4 | 21 | 0.6000 |
| | | Superdome concrete | – | Superdome clay | 3 | 4 | 9 | 1 |
| | 2 months | Small concrete rod | – | Large concrete rod | 6 | 6 | 24 | 1 |
| | | Small concrete rod | – | Small clay rod | 6 | 6 | 22 | 1 |
| | | Small concrete rod | – | Large clay rod | 6 | 6 | 17 | 1 |
| | | Small concrete rod | – | Superdome concrete | 6 | 3 | 12 | 1 |
| | | Small concrete rod | – | Superdome clay | 6 | 4 | 11 | 1 |
| | | Large concrete rod | – | Small clay rod | 6 | 6 | 13 | 1 |
| | | Large concrete rod | – | Large clay rod | 6 | 6 | 11 | 1 |
| | | Large concrete rod | – | Superdome concrete | 6 | 3 | 8 | 1 |
| | | Large concrete rod | – | Superdome clay | 6 | 4 | 7 | 1 |
| | | Small clay rod | – | Large clay rod | 6 | 6 | 15 | 1 |
| | | Small clay rod | – | Superdome concrete | 6 | 3 | 12 | 1 |
| | | Small clay rod | – | Superdome clay | 6 | 4 | 8 | 1 |
| | | Large clay rod | – | Superdome concrete | 6 | 3 | 12 | 1 |
| | | Large clay rod | – | Superdome clay | 6 | 4 | 8 | 1 |
| | | Superdome concrete | – | Superdome clay | 3 | 4 | 3 | 1 |
| | 6 months | Small concrete rod | – | Large concrete rod | 6 | 6 | 14 | 1 |
| | | Small concrete rod | – | Small clay rod | 6 | 6 | 12 | 1 |
| | | Small concrete rod | – | Large clay rod | 6 | 6 | 0 | 0.0740 |
| | | Small concrete rod | – | Superdome concrete | 6 | 3 | 1 | 0.4580 |
| | | Small concrete rod | – | Superdome clay | 6 | 4 | 2 | 0.4580 |

| | | | | | | | | |
|---|---|---|---|---|---|---|---|---|
| | | Large concrete rod | – | Small clay rod | 6 | 6 | 16 | 1 |
| | | Large concrete rod | – | Large clay rod | 6 | 6 | 0 | 0.0740 |
| | | Large concrete rod | – | Superdome concrete | 6 | 3 | 0 | 0.3300 |
| | | Large concrete rod | – | Superdome clay | 6 | 4 | 2 | 0.4580 |
| | | Small clay rod | – | Large clay rod | 6 | 6 | 1 | 0.1040 |
| | | Small clay rod | – | Superdome concrete | 6 | 3 | 2 | 0.7250 |
| | | Small clay rod | – | Superdome clay | 6 | 4 | 5 | 0.9780 |
| | | Large clay rod | – | Superdome concrete | 6 | 3 | 16 | 0.7250 |
| | | Large clay rod | – | Superdome clay | 6 | 4 | 19 | 0.9780 |
| | | Superdome concrete | – | Superdome clay | 3 | 4 | 7 | 1 |
| Planar recruit density | 1 week | Small concrete rod | – | Large concrete rod | 6 | 6 | 36 | **0.0320** |
| | | Small concrete rod | – | Small clay rod | 6 | 6 | 36 | **0.0320** |
| | | Small concrete rod | – | Large clay rod | 6 | 6 | 36 | **0.0320** |
| | | Small concrete rod | – | Superdome concrete | 6 | 3 | 18 | 0.1430 |
| | | Small concrete rod | – | Superdome clay | 6 | 4 | 24 | 0.0950 |
| | | Large concrete rod | – | Small clay rod | 6 | 6 | 11 | 0.6200 |
| | | Large concrete rod | – | Large clay rod | 6 | 6 | 35 | **0.0480** |
| | | Large concrete rod | – | Superdome concrete | 6 | 3 | 18 | 0.1430 |
| | | Large concrete rod | – | Superdome clay | 6 | 4 | 24 | 0.0950 |
| | | Small clay rod | – | Large clay rod | 6 | 6 | 36 | **0.0320** |
| | | Small clay rod | – | Superdome concrete | 6 | 3 | 18 | 0.1430 |
| | | Small clay rod | – | Superdome clay | 6 | 4 | 24 | 0.0950 |
| | | Large clay rod | – | Superdome concrete | 6 | 3 | 18 | 0.1430 |
| | | Large clay rod | – | Superdome clay | 6 | 4 | 24 | 0.0950 |
| | | Superdome concrete | – | Superdome clay | 3 | 4 | 9 | 0.6200 |
| | 2 months | Small concrete rod | – | Large concrete rod | 6 | 6 | 36 | 0.0750 |
| | | Small concrete rod | – | Small clay rod | 6 | 6 | 23.5 | 0.7440 |
| | | Small concrete rod | – | Large clay rod | 6 | 6 | 36 | 0.0750 |
| | | Small concrete rod | – | Superdome concrete | 6 | 3 | 18 | 0.1820 |
| | | Small concrete rod | – | Superdome clay | 6 | 4 | 24 | 0.1250 |
| | | Large concrete rod | – | Small clay rod | 6 | 6 | 0 | 0.0750 |
| | | Large concrete rod | – | Large clay rod | 6 | 6 | 4 | 0.1820 |
| | | Large concrete rod | – | Superdome concrete | 6 | 3 | 18 | 0.1820 |
| | | Large concrete rod | – | Superdome clay | 6 | 4 | 24 | 0.1050 |
| | | Small clay rod | – | Large clay rod | 6 | 6 | 36 | 0.0750 |
| | | Small clay rod | – | Superdome concrete | 6 | 3 | 18 | 0.1820 |
| | | Small clay rod | – | Superdome clay | 6 | 4 | 24 | 0.1250 |
| | | Large clay rod | – | Superdome concrete | 6 | 3 | 18 | 0.1820 |
| | | Large clay rod | – | Superdome clay | 6 | 4 | 24 | 0.1050 |
| | | Superdome concrete | – | Superdome clay | 3 | 4 | 3 | 0.7440 |
| | 6 months | Small concrete rod | – | Large concrete rod | 6 | 6 | 36 | 0.0740 |
| | | Small concrete rod | – | Small clay rod | 6 | 6 | 17 | 1 |
| | | Small concrete rod | – | Large clay rod | 6 | 6 | 10 | 0.9080 |
| | | Small concrete rod | – | Superdome concrete | 6 | 3 | 18 | 0.2100 |
| | | Small concrete rod | – | Superdome clay | 6 | 4 | 24 | 0.1680 |

| | | | | | | |
|---|---|---|---|---|---|---|
| Large concrete rod | – | Small clay rod | 6 | 6 | 2 | 0.1680 |
| Large concrete rod | – | Large clay rod | 6 | 6 | 0 | 0.0740 |
| Large concrete rod | – | Superdome concrete | 6 | 3 | 18 | 0.2100 |
| Large concrete rod | – | Superdome clay | 6 | 4 | 24 | 0.1680 |
| Small clay rod | – | Large clay rod | 6 | 6 | 11 | 0.9080 |
| Small clay rod | – | Superdome concrete | 6 | 3 | 18 | 0.2100 |
| Small clay rod | – | Superdome clay | 6 | 4 | 24 | 0.1680 |
| Large clay rod | – | Superdome concrete | 6 | 3 | 18 | 0.2100 |
| Large clay rod | – | Superdome clay | 6 | 4 | 24 | 0.1680 |
| Superdome concrete | – | Superdome clay | 3 | 4 | 7 | 1 |

**Table S3**: Differences in recruit densities (recruits per $cm^2$) between the outside (exposed) and inside (recesses) areas of the six different shapes (superdome clay, superdome concrete, large clay rod, small clay rod, large concrete rod, small concrete rod). Results are derived from Wilcoxon rank-sum tests. Number of replicates per test group (n1 and n2), test statistic (*W*), and p-value are given with bold values indicating significance ($p < 0.05$).

| Module type | Timepoint | Contrast | | | n1 | n2 | statistic | p |
|---|---|---|---|---|---|---|---|---|
| Small concrete rod | 1 week | Outside | – | Inside | 6 | 6 | 0 | **0.0050** |
| Small concrete rod | 2 months | Outside | – | Inside | 6 | 6 | 0 | **0.0041** |
| Small concrete rod | 6 months | Outside | – | Inside | 6 | 6 | 0 | **0.0042** |
| Large concrete rod | 1 week | Outside | – | Inside | 6 | 6 | 0 | **0.0022** |
| Large concrete rod | 2 months | Outside | – | Inside | 6 | 6 | 0 | **0.0047** |
| Large concrete rod | 6 months | Outside | – | Inside | 6 | 6 | 0 | **0.0043** |
| Small clay rod | 1 week | Outside | – | Inside | 6 | 6 | 0 | **0.0050** |
| Small clay rod | 2 months | Outside | – | Inside | 6 | 6 | 0 | **0.0048** |
| Small clay rod | 6 months | Outside | – | Inside | 6 | 6 | 2 | **0.0121** |
| Large clay rod | 1 week | Outside | – | Inside | 6 | 6 | 0 | **0.0022** |
| Large clay rod | 2 months | Outside | – | Inside | 6 | 6 | 0 | **0.0049** |
| Large clay rod | 6 months | Outside | – | Inside | 6 | 6 | 0 | **0.0049** |
| Superdome concrete | 1 week | Outside | – | Inside | 3 | 3 | 0 | 0.1000 |
| Superdome concrete | 2 months | Outside | – | Inside | 3 | 3 | 0 | 0.0593 |
| Superdome concrete | 6 months | Outside | – | Inside | 3 | 3 | 0 | 0.0593 |
| Superdome clay | 1 week | Outside | – | Inside | 4 | 4 | 0 | **0.0294** |
| Superdome clay | 2 months | Outside | – | Inside | 4 | 4 | 0 | **0.0211** |
| Superdome clay | 6 months | Outside | – | Inside | 4 | 4 | 0 | **0.0202** |

**Table S4**: Differences in survival rate of recruits (%) between the six different shapes (superdome clay, superdome concrete, large clay rod, small clay rod, large concrete rod, small concrete rod) from 1 week to 2 months, and to six months. Results are derived from Wilcoxon rank-sum tests followed by Holm adjustment for multiple testing. Number of replicates per test group (n1 and n2), test statistic (*W*), and p-value are given with bold values indicating significance ($p < 0.05$).

| Change period | Group 1 | | Group 2 | n1 | n2 | statistic | p |
|---|---|---|---|---|---|---|---|
| 1 week to 2 months | Small concrete rod | – | Large concrete rod | 6 | 6 | 23 | 1 |
| | Small concrete rod | – | Small clay rod | 6 | 6 | 1 | 0.0650 |
| | Small concrete rod | – | Large clay rod | 6 | 6 | 2 | 0.1210 |
| | Small concrete rod | – | Superdome concrete | 6 | 3 | 8 | 1 |
| | Small concrete rod | – | Superdome clay | 6 | 4 | 0 | 0.1210 |
| | Large concrete rod | – | Small clay rod | 6 | 6 | 2 | 0.1210 |
| | Large concrete rod | – | Large clay rod | 6 | 6 | 2 | 0.1210 |
| | Large concrete rod | – | Superdome concrete | 6 | 3 | 5 | 1 |
| | Large concrete rod | – | Superdome clay | 6 | 4 | 0 | 0.1210 |
| | Small clay rod | – | Large clay rod | 6 | 6 | 17 | 1 |
| | Small clay rod | – | Superdome concrete | 6 | 3 | 14 | 1 |
| | Small clay rod | – | Superdome clay | 6 | 4 | 3 | 0.6000 |
| | Large clay rod | – | Superdome concrete | 6 | 3 | 14 | 1 |
| | Large clay rod | – | Superdome clay | 6 | 4 | 3 | 0.6000 |
| | Superdome concrete | – | Superdome clay | 3 | 4 | 1 | 0.7980 |
| 2 months to 6 months | Small concrete rod | – | Large concrete rod | 6 | 6 | 11 | 1 |
| | Small concrete rod | – | Small clay rod | 6 | 6 | 14 | 1 |
| | Small concrete rod | – | Large clay rod | 6 | 6 | 0 | **0.0320** |
| | Small concrete rod | – | Superdome concrete | 6 | 3 | 0 | 0.2860 |
| | Small concrete rod | – | Superdome clay | 6 | 4 | 3 | 0.6000 |
| | Large concrete rod | – | Small clay rod | 6 | 6 | 20 | 1 |
| | Large concrete rod | – | Large clay rod | 6 | 6 | 0 | **0.0320** |
| | Large concrete rod | – | Superdome concrete | 6 | 3 | 1 | 0.4760 |
| | Large concrete rod | – | Superdome clay | 6 | 4 | 7 | 1 |
| | Small clay rod | – | Large clay rod | 6 | 6 | 0 | **0.0320** |
| | Small clay rod | – | Superdome concrete | 6 | 3 | 0 | 0.2860 |
| | Small clay rod | – | Superdome clay | 6 | 4 | 5 | 1 |
| | Large clay rod | – | Superdome concrete | 6 | 3 | 16 | 0.7620 |
| | Large clay rod | – | Superdome clay | 6 | 4 | 20 | 0.7980 |
| | Superdome concrete | – | Superdome clay | 3 | 4 | 8 | 1 |

**Table S5:** Effects of timepoint, module type, and their interaction on coral recruit density (recruits per $cm^2$), assessed using a Tweedie generalized linear model with a log link function. Likelihood-ratio tests ($\chi^2$) were used to evaluate the significance of each model term. Values in bold indicate statistically significant effects ($p < 0.05$).

| Effect | Chisq | df | p |
|---|---|---|---|
| Timepoint | 3244.13825 | 2 | **<0.0001** |
| Module type | 16.5078648 | 5 | **0.0055** |
| Timepoint x Module type | 130.233398 | 10 | **<0.0001** |

**Table S6**: Differences in recruit densities (recruits per $cm^2$) between the six different shapes (superdome clay, superdome concrete, large clay rod, small clay rod, large concrete rod, small concrete rod) and natural substrate in the barrier reef after two months. Results are derived from Wilcoxon rank-sum tests followed by Holm adjustment for multiple testing. Number of replicates per test group (n1 and n2), test statistic (*W*), and p-value are given with bold values indicating significance ($p < 0.05$).

| Contrast | | | n1 | n2 | Statistic | p |
|---|---|---|---|---|---|---|
| Reef | – | Superdome clay | 29 | 4 | 0 | **0.0020** |
| Reef | – | Superdome concrete | 29 | 3 | 0 | **0.0050** |
| Reef | – | Large clay rod | 29 | 6 | 0 | **0.0009** |
| Reef | – | Small clay rod | 29 | 6 | 0 | **0.0009** |
| Reef | – | Large concrete rod | 29 | 6 | 0 | **0.0009** |
| Reef | – | Small concrete rod | 29 | 6 | 0 | **0.0009** |

**Table S7**: List of R3D consortium members. Lab leads are noted in bold.

| Author | Institution | ORCID |
|---|---|---|
| **Benjamin A. Jones** | Applied Research Laboratory at the University of Hawai‘i | 0009-0000-2692-7443 |
| Joshua Levy | Applied Research Laboratory at the University of Hawai‘i | |
| Sean Mahaffey | Applied Research Laboratory at the University of Hawai‘i | |
| Aricia Argyris | Applied Research Laboratory at the University of Hawai‘i | |
| Mark Aruda | Applied Research Laboratory at the University of Hawai‘i | |
| Ian Robertson | Applied Research Laboratory at the University of Hawai‘i | |
| **Zhenhua Huang** | University of Hawaii at Manoa | 0000-0001-6665-7230 |
| Ayrton Medina-Rodriguez | University of Hawaii at Manoa | 0000-0002-0666-9472 |
| Mert Gokdepe | University of Hawaii at Manoa | |
| Brady Halvorson | University of Hawaii at Manoa | |
| Jon Chase | University of Hawaii at Manoa | |
| Charlotte White | University of Hawaii at Manoa | |
| Cami Dillon | University of Hawaii at Manoa | |
| Kristian McDonald | University of Hawaii at Manoa | |
| Anna Mikkelsen | University of Hawaii at Manoa | |
| **Josh Madin** | Hawai‘i Institute of Marine Biology | 0000-0002-5005-6227 |
| Mollie Asbury | Hawai‘i Institute of Marine Biology | |
| Jessica Reichert | Hawai‘i Institute of Marine Biology | 0000-0003-2245-4188 |
| Hendrikje Jorissen | Hawai‘i Institute of Marine Biology | |
| Nina Schiettekatte | Hawai‘i Institute of Marine Biology | |
| Marion Chapeau | Hawai‘i Institute of Marine Biology | |
| **Rob Toonen** | Hawai‘i Institute of Marine Biology | 0000-0001-6339-4340 |
| Christopher R. Suchocki | Hawai‘i Institute of Marine Biology | 0000-0001-6811-1987 |
| Van Wishingrad | Hawai‘i Institute of Marine Biology | 0000-0002-6256-4018 |
| Chris Jury | Hawai‘i Institute of Marine Biology | |
| Daniel Schar | Hawai‘i Institute of Marine Biology | |
| Madeleine Hardt | Hawai‘i Institute of Marine Biology | |
| Claire Lewis | Hawai‘i Institute of Marine Biology | 0000-0003-1081-2734 |
| Claire Bardin | Hawai‘i Institute of Marine Biology | |
| Joshua Kualani | Hawai‘i Institute of Marine Biology | |
| **Crawford Drury** | Hawai‘i Institute of Marine Biology | 0000-0001-8853-416X |
| **Kira Hughes** | Hawai‘i Institute of Marine Biology | |
| Josh Hancock | Hawai‘i Institute of Marine Biology | 0000-0002-4814-1385 |
| Carlo Caruso | Hawai‘i Institute of Marine Biology | |
| **Andrea Grottoli** | Ohio State University | 0000-0001-6053-9452 |
| Shannon Dixon | Ohio State University | 0009-0007-9882-7936 |
| Ann Marie Hulver | Ohio State University | 0000-0003-0466-9070 |
| **Joshua D. Voss** | Florida Atlantic University | 0000-0002-0653-2767 |
| Allison Klein | Florida Atlantic University | 0009-0004-2670-4222 |
| **Siddhartha Verma** | Florida Atlantic University | 0000-0002-8941-0633 |
| Alejandro Alvaro | Florida Atlantic University | |
| **Richard Argall** | Makai Ocean Engineering | |

| | |
|---|---|
| Kevin Chun | Makai Ocean Engineering |
| William Hicks | Makai Ocean Engineering |
| Alex LeBon | Makai Ocean Engineering |
| John Yeh | Makai Ocean Engineering |
| **Aaron Thode** | Scripps Institution of Oceanography, UC San Diego |
| Oceane Boulais | Scripps Institution of Oceanography, UC San Diego |
| **Daniel Wangpraseurt** | Scripps Institution of Oceanography, UC San Diego |
| Samapti Kundu | Scripps Institution of Oceanography, UC San Diego |
| Natalie Levy | Scripps Institution of Oceanography, UC San Diego |
| Lindsey Badder | Scripps Institution of Oceanography, UC San Diego |
| Stefan Kolle | Scripps Institution of Oceanography, UC San Diego |
